# Delayed protective effect of chronic variable stress on optic tract axonal degeneration after experimental TBI

**DOI:** 10.1101/2025.10.02.680140

**Authors:** Macy Urig, Jordyn N. Torrens, Shelby M. Hetzer, Emma G. Burke, John M. Riccobono, Abby Atreya, Samantha Lingo, James P. Herman, Nathan K. Evanson

## Abstract

Of the 2.8 million individuals who seek medical attention for traumatic brain injury (TBI) each year, nearly 300,000 require hospitalization, with up to 60% of these needing intensive care. Intensive care treatment of TBI involves stressful events such as sleep disruption, noise, and painful procedures, potentially leading to chronic stress in patients undergoing such treatment. Given that physiologic stress can exacerbate neuroinflammation and impair normal neural function, we hypothesized that chronic variable stress (CVS) following TBI would exacerbate behavioral and pathological outcomes. We tested this hypothesis by subjecting adolescent male mice to blunt TBI, followed by two weeks of CVS or control conditions. We assessed brain pathologic responses to injury 2-, 5-, 20-, and 28-weeks post-injury. We found chronic optic tract degeneration by Fluoro-jade B staining in TBI groups. Unexpectedly, CVS+TBI mice did not show evidence of optic tract axon degeneration 20 weeks after injury, but did at the other time points. CVS led to increased microglial phagocytic markers early after injury, regardless of TBI status, and TBI led to increased microglial phagocytic markers in a delayed fashion as well. Notably, microglial phagocytosis markers were not elevated in TBI+CVS groups compared to TBI only groups 20 weeks post-injury. There was no effect of TBI or CVS on behavioral measures taken at the end of CVS. These findings suggest a delayed, but not permanent, protective effect on axonal degeneration after TBI, potentially related to altered microglial and astrocytic phagocytic activity.

## Introduction

Nearly 300,000 of the 2.8 million patients who seek medical assistance after a traumatic brain injury (TBI) each year will be hospitalized, with up to 60% requiring admission to an intensive care unit (Levant et al., 2016; Taylor et al., 2017). While in intensive care, patients are exposed to many stressors such as pain, uncomfortable procedures and medical equipment, loud noises, and disrupted sleep (Krampe et al., 2021). Intensive care patients report high incidence of significant stressors regardless of type of intensive care unit (de Sá Dias et al., 2015). In addition to the disruptive nature of initial hospitalization, TBI also alters physiologic stress responses, including both acute and chronic hypothalamus-pituitary-adrenal (HPA) axis stress responses (Taylor et al., 2006; Taylor et al., 2008).

The intersection of stress and injury is especially relevant to adolescents and young adults. Adolescence is a time of both increased risk for TBI and also altered physiologic responses to stress, which can have lasting effects on brain development (Eiland and Romeo, 2013). Importantly, chronic stress during adolescence can lead to distinct long-term outcomes when compared to chronic stress in adulthood (Cotella et al., 2019). Changes in physiologic stress reactivity can have significant implications for long-term outcomes of TBI (Weil and Karelina, 2019). Animal models of adolescent TBI have shown increased hippocampal neurodegeneration related to traumatic brain injury when compared to adult models (Guilhaume-Correa et al., 2020). From the intersection of these findings, it is clear that the interaction of chronic stress and brain injury in adolescence represents an important relationship to understand.

We previously described long-lasting axonal degeneration in the optic tracts of mice after experimental weight-drop TBI in adolescence (Hetzer et al., 2021a). This degeneration is associated with neuroinflammation and gliosis in optic tracts and major projection targets of the optic nerve. Because chronic stress can alter neuroinflammation (White et al., 2024), and because of the above-noted chronic alterations in physiologic stress responses resulting from TBI and the associated medical care, we hypothesized that chronic stress post-injury would exacerbate behavioral and pathological outcomes after experimental TBI. To test this hypothesis, we compared behavioral and histologic changes between adolescent male mice with TBI exposed to chronic variable stress (CVS) and those not exposed to CVS post-injury.

## Methods

### Animals

Experiments were performed using 6-week-old adolescent male C57BL/6J mice (Jackson Laboratories, Bar Harbor, ME). Mice were housed under a 14h:10h light:dark schedule in pressurized, individually ventilated cage racks, with 4 mice per cage, and were given ad libitum access to water and standard rodent chow. Animals habituated to the vivarium for one week prior to undergoing TBI and subsequent procedures. The University of Cincinnati Institutional Animal Care and Use Committee approved all experimental procedures.

### Traumatic Brain Injury

Closed-head injury was performed by weight drop, as previously described (Yang et al., 2013; Evanson et al., 2018). Briefly, mice were anesthetized using isoflurane (2-3%), then placed in prone position under a 400 g metal rod raised 1.5 cm above the mouse’s head. The rod was dropped onto the intact scalp roughly above bregma. After TBI, mice were placed under an oxygen hood made from a pipet box connected to an oxygen concentrator and were exposed to 100% oxygen until normal breathing was regained. Mice were observed for recovery of righting reflex before being returned to their home cages. Injured mice will hereon be referred to as TBI mice. Non-injured, sham animals were anesthetized, weighed, and allowed to right before being returned to their home cages. All TBI procedures for this experiment were performed on the same day in random order by cage, using a random number generator. One cohort of mice was used for behavioral testing as described below. Four other cohorts were used for histologic studies performed at 2-weeks, 5-weeks, 20-weeks, or 28-weeks post injury. Figure 1A outlines the experimental procedures used.

**Figure 1.**
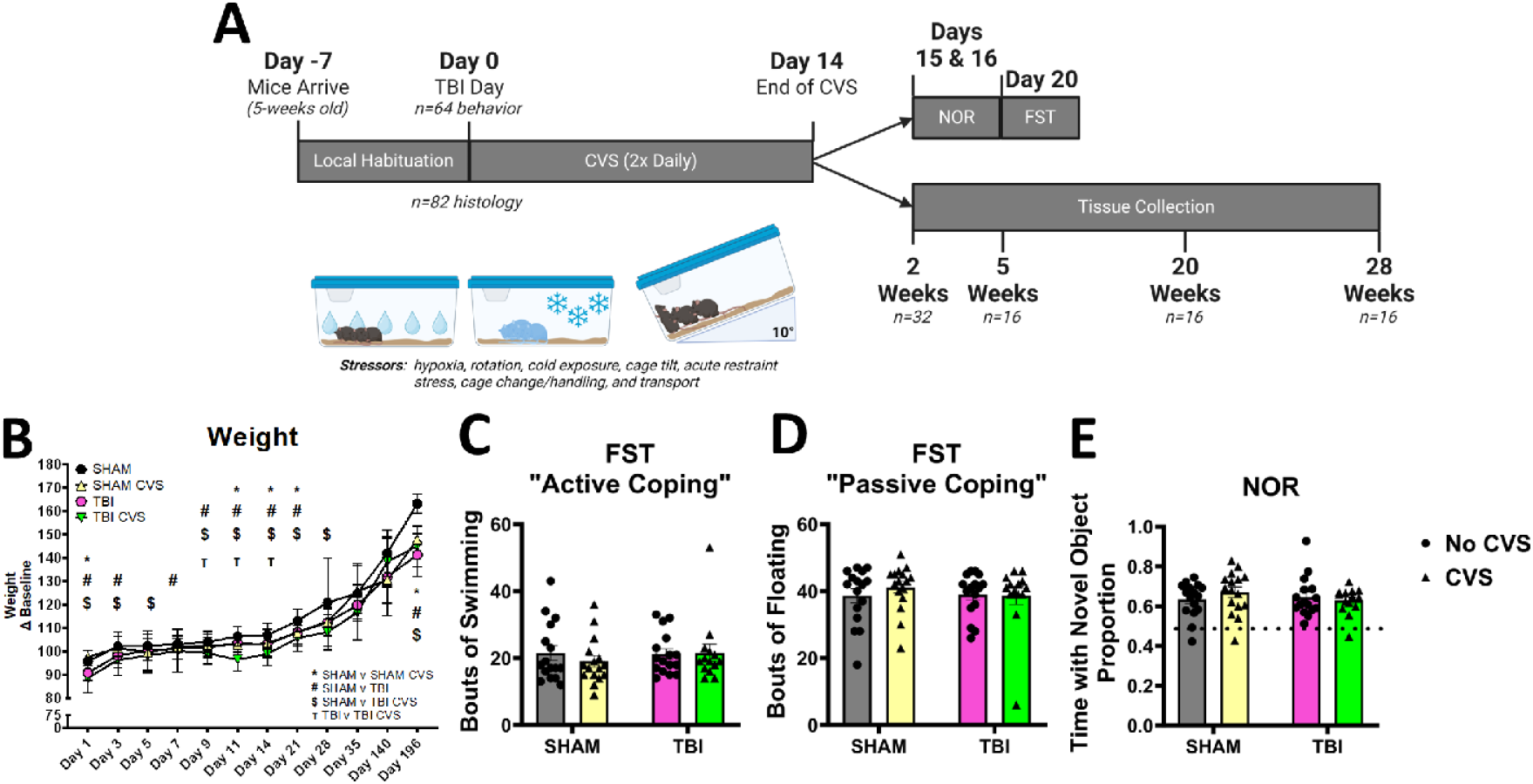
Experimental Timeline, weight, and behavior. A) Timeline of experimental procedures, showing behavior and histology cohorts, and brief outline of CVS procedures. B) Weight trend of mice over the course of experimental procedures, with % of baseline weight on the y axis. Statistical significance (p < 0.05) is indicated by symbols as outlined in the figure. C) FST active coping (swimming behavior). D) FST passive coping (immobility). E) NOR proportion of time with novel object. There was no significant difference between groups in active or passive coping during FST or in NOR performance.

### Chronic Variable Stress

Mice were divided into groups that were exposed to a chronic variable stress (CVS) paradigm, adapted from (Herman et al., 1995). Briefly, mice were exposed to twice daily stressors (AM between 9:00–12:00 and PM between 2:00–6:00) over a two week-period. Stressors were randomly assigned with no duplicated stressors within the same day. Stressors included hypoxia (30 min with 8% O_2_ and 92%N); rotation (cage placed on orbital shaker at 100 rpm for 1 h), cold exposure (4 °C for 30 minutes with no food/water or bedding); cage tilt (10° angle overnight), acute restraint stress (60 minutes in a 50mL tube with nose and tail holes cut out), cage change, transport (30 minutes walking with cages placed on a wheeled cart), or wet bedding (AM stressor 9:00-12:00PM). During the two weeks of CVS, control animals were maintained under standard animal husbandry conditions.

### Forced Swim Test (FST)

FST was performed on post-injury day 20, as previously described (Solomon et al., 2015). Briefly, mice were individually placed in a 2-liter beaker filled to 1200 ml with water at 23 +/- 2 °C and videorecorded for 10 minutes. Behavior was scored by blinded observers who observed behaviors at 5 second intervals and scored them for whether they were swimming or floating at each interval. The total number of swimming or floating instances were tallied for each mouse.

### Novel Object Recognition (NOR)

NOR was performed on post-injury days 15 and 16, as we previously described (Guilhaume-Correa et al., 2020). Briefly, on the first day of testing a mouse was placed in a guinea pig cage and allowed to explore two identical objects for 10 minutes. After this time, the mouse was returned to its home cage. On the second day of testing, one of the objects was replaced by a novel object, and mice were placed in the cage and allowed to explore the objects for 5 minutes. Behavior was videorecorded and scored by blinded investigators for the amount of time spent exploring each object. Results are expressed as the proportion of time spent exploring the novel object out of the total exploration time of both objects.

### Histology

#### Tissue Collection

We removed brains for histologic and immunohistochemical analyses as previously described (Hetzer et al., 2021b). Briefly, mice were euthanized by barbiturate overdose, then perfused with 4% paraformaldehyde in phosphate buffered saline.

Brains were collected and post-fixed in 4% paraformaldehyde overnight then immersed in 30% sucrose. Sucrose-impregnated brains were sectioned into 30 micron sections using a freezing stage sliding microtome (Leica, Bannockburn, IL). Sections were stored at -20°C in cryoprotective solution (0.01M phosphate-buffered saline, 1% polyvinyl-pyrrolidone (Sigma-Aldrich, St. Louis, MO; Cat# PVP-40), 30% ethylene glycol (Thermo-Fisher, Waltham, MA; Cat# E178-4), and 30% sucrose (Thermo-Fisher; Cat# S5-3)).

#### Fluoro-jade B Staining

Fluoro-jade B (FJ-B; Histo-Chem, Jackson, AR; CAT# 1FJB), a marker for degenerating neurons and axons (Schmued and Hopkins, 2000), was used to stain tissue sections according to the manufacturer’s directions as we previously reported (Guilhaume-Correa et al., 2020), with slight modifications (sections were incubated in 0.06% potassium permanganate for 5 minutes, then submerged in 0.0001% FJ-B solution for 5 minutes to reduce background fluorescence). After staining, slides were air dried in the dark. Slides were stored without coverslips in a slide box and imaged immediately after drying, also to reduce background fluorescence.

#### Immunohistochemistry

Free-floating brain sections were blocked for one hour at room temperature (1% bovine serum albumin, 5% normal goat serum, 0.2% Triton X-100) before incubation in primary antibodies against LAMP2 (1:500; (Millipore Cat# MABC40, RRID:AB_10808497), glial fibrillary acidic protein (GFAP, 1:3000; DAKO, Santa Clara, CA; CAT# Z0334; RRID AB_10013382), ionized calcium-binding adaptor molecule 1 (Iba-1, 1:1000; Wako cat# 019-19741, RRID:AB_839504), or Cluster of differentiation factor 68 (CD68,1:750, Bio-Rad Cat# MCA1957A488T, RRID:AB_1102282) overnight at 4°C. For double labeling, the second primary antibody was then incubated overnight at 4°C. On the third day, tissue was incubated with goat-anti-rabbit Alexa Fluor 647 and goat anti-rat Alexa Fluor 546 antibodies (Jackson Immunochemicals, West Grove, PA; cat# 711-165-152, RRID AB_2307443) at 1:500 dilution overnight at 4°C. Slides were cover slipped using antifading polyvinyl alcohol mounting medium (Sigma-Aldrich, St.Louis, MO, Cat# 10981).

#### Image Analysis

For FJ-B, images of left and right regions of interest were captured on a Nikon C2 Plus Confocal Microscope (Nikon Corporation, Melville, New York) using 20X objective lens. Imaged regions included the optic tract (OT), paraventricular nucleus of the hypothalamus (PVN), thalamic lateral geniculate nucleus (LGN), and superior colliculus (SC). A blinded investigator took all pictures. FJ-B positive area was measured using Nikon Elements Analysis software (Nikon) as we previously described (Hetzer et al., 2023).

For GFAP/LAMP2 and IBA-1/CD68 double stained sections, images of left and right OT and PVN were captured on a Nikon C2 Plus Confocal Microscope (Nikon) using a 40X objective lens. Volume of GFAP, LAMP2, Iba-1, and CD68 were calculated using the 3D measurement tools and summating each positively stained object with thresholds and imaging parameters held constant across images in Nikon Elements (Nikon).

#### Statistical Analyses

Weight change was analyzed using a mixed effects model with injury x stress x day as independent variables, using GraphPad Prism 9 (GraphPad, San Diego, CA). Immunofluorescence results were analyzed using 2-way ANOVA (injury x stress as independent variables, with each time point being analyzed independently) using GraphPad Prism 9 (GraphPad). Critical alpha was set a priori at p < 0.05.

## Results

### Weight and Behavioral Effects of TBI and CVS

Weights were recorded every 2-3 days until 14 days post injury (the final day of CVS), and just prior to euthanasia (Figure 1B). There was a main effect of both injury and stress with significant stress x time and injury x time interactions. Stress effects were significant between sham and sham + CVS mice starting on day 11 and lasting throughout the study. Moreover, TBI + CVS mice weighed the least out of the four groups from day 11 onward. For a detailed list of relevant statistical tests reflected in Figure 1B, see Table S1.

We assessed mice for changes in coping behavior during FST. One week after the end of CVS, mice were recorded and behavior scored for active coping behaviors (i.e., swimming) and passive coping behaviors (i.e., floating). We found no differences between any groups for active or passive coping behaviors (Figure 1C,D). We also found no differences in NOR performance between any groups (Figure 1E).

### Fluoro-jade B Neurodegeneration Stain

In the optic tract, there was no positive FJ-B staining in sham groups at any time point (Figure 2). Two-weeks post injury, FJ-B was significantly higher for both the TBI and TBI+CVS groups compared to uninjured mice (main effect of injury p=0.0001, Figure 2B). This increase was also present in the 5-week cohort (main effect of injury p=0.0005, Figure 2C). In the 20-week cohort, there was a significant interaction between injury and stress (p=0.002); the TBI+CVS group had no positive staining (Figure 2D). At 28-weeks there was a significant main effect of injury (p=0.005), but post hoc tests were not significant for either injured group compared to their respective sham group. Subjectively, there was still positive FJ-B staining in the optic tract of these mice (Figure 2A). Similar results were reflected in optic nerve projection targets including the LGN and SC (Figure S1, Table S2). In the PVN, there was no positive FJ-B at any time point in any group (Figure 3). For detailed statistical results from FJ-B staining in the optic tracts, see Table 1.

**Table 1.** Statistical Results 2-Way ANOVA – FJ-B staining.

| Table 1. Statistical Results 2-Way ANOVA – FJ-B staining |  |  |  |  |  |
| --- | --- | --- | --- | --- | --- |
| Timepoint | Comparison | Optic Tract |  | PVN |  |
|  |  | Test Result | p Value | Test Result | p Value |
| 2 weeks | Injury | $F_{1,25} = 27.46$ | **0.0001** | $F_{1,25} = 0.79$ | 0.38 |
| | Stress | $F_{1,25} = 0.16$ | 0.69 | $F_{1,25} = 0.15$ | 0.71 |
| | Interaction | $F_{1,25} = 0.89$ | 0.35 | $F_{1,25} = 0.86$ | 0.36 |
| 5 weeks | Main effect of injury | $F_{1,12} = 22.78$ | **0.0005** | $F_{1,12} = 0.99$ | 0.26 |
| 5 weeks | Main effect of stress | $F_{1,12} = 0.003$ | 0.96 | $F_{1,12} = 0.12$ | 0.74 |
| 5 weeks | Interaction | $F_{1,12} = 0.0005$ | 0.98 | $F_{1,12} = 1.41$ | 0.34 |
| 20 weeks | Main effect of injury | $F_{1,12} = 15.31$ | **0.002** | $F_{1,12} = 1.57$ | 0.23 |
| 20 weeks | Main effect of stress | $F_{1,12} = 12.98$ | **0.004** | $F_{1,12} = 0.53$ | 0.48 |
| 20 weeks | Interaction | $F_{1,12} = 14.77$ | **0.002** | $F_{1,12} = 0.72$ | 0.41 |
| 28 weeks | Main effect of injury | $F_{1,12} = 11.75$ | **0.005** | $F_{1,11} = 0.50$ | 0.50 |
| 28 weeks | Main effect of stress | $F_{1,12} = 0.10$ | 0.76 | $F_{1,11} = 0.07$ | 0.80 |
| 28 weeks | Interaction | $F_{1,12} = 0.003$ | 0.96 | $F_{1,11} = 4.00$ | 0.07 |

**Figure 2:**
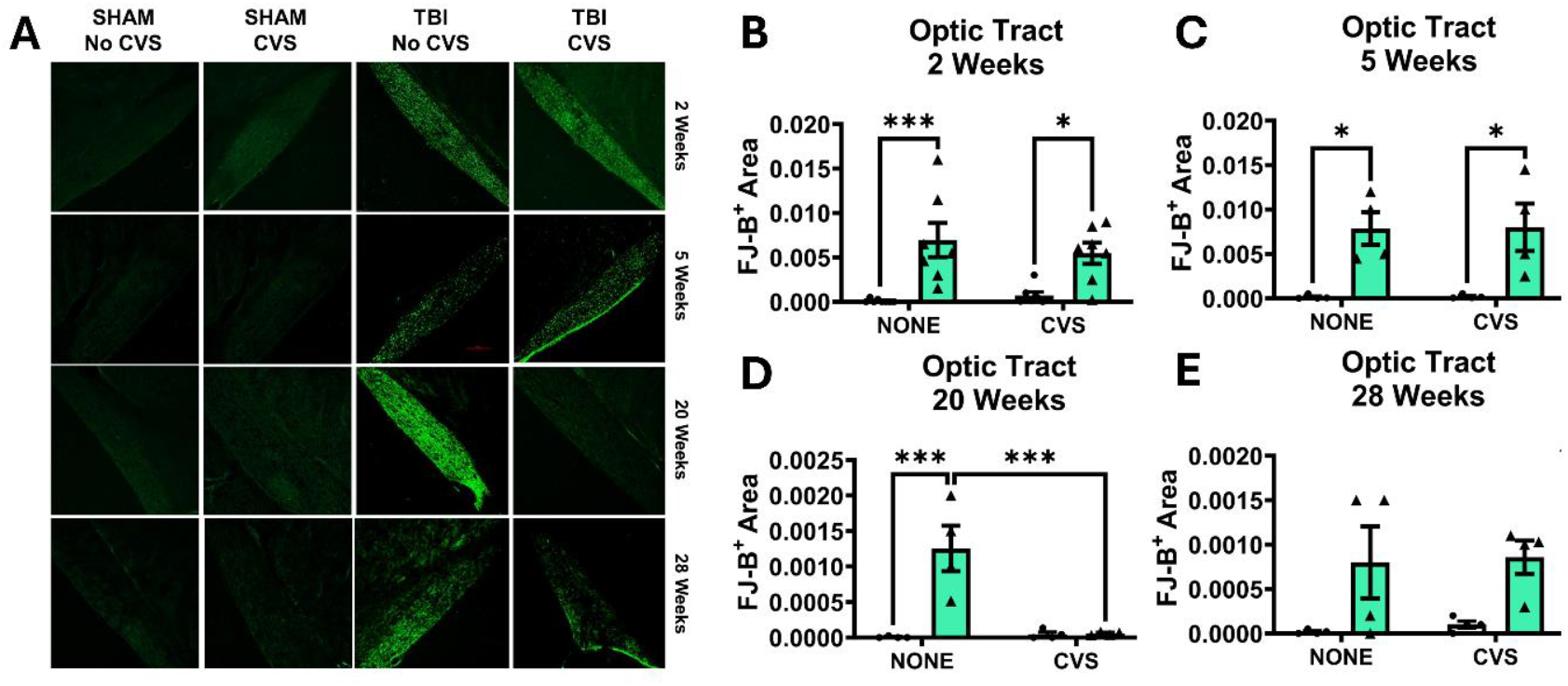
Axonal degeneration in optic tract by FJ-B staining. A) Representative images from FJ-B staining, in optic tract, using 20X objective. Also shown is quantified FJ-B staining in optic tract at B) 2 weeks, C) 5 weeks, D) 20 weeks, and E) 28 weeks after injury. Green shaded bars represent TBI groups in all graphs. * p < 0.05; *** p < 0.001

**Figure 3:**
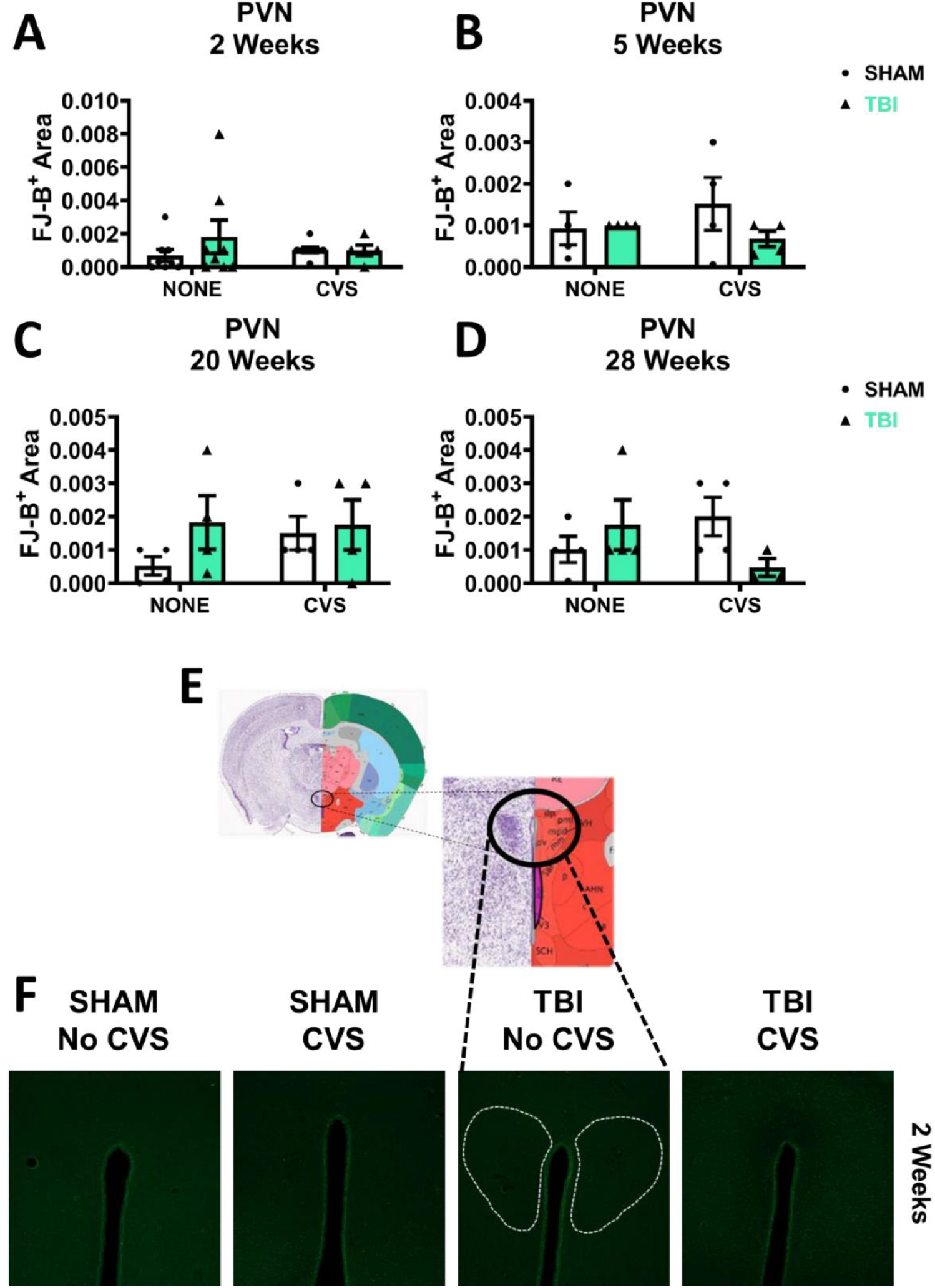
There was no significant axonal or neuronal degeneration seen in the PVN by FJ-B staining. FJ-B staining is shown in the PVN at A) 2 weeks, B) 5 weeks, C) 20 weeks, and D) 28 weeks after injury. E) Diagram of the location of PVN where photomicrographs were taken. F) Representative images for FJ-B staining in the PVN, taken using 10X objective lens.

### Immunohistochemistry

We immunostained brain slices with antibodies against Iba-1 and CD68 to assess their overlap as a marker of microglial activation/phagocytosis in the OT and PVN (Figure 4, Figure 5, Table 2). For measures of CD68 volume in the OT, there was a main effect of CVS at 2 weeks, a main effect of TBI at 5 weeks, and a stress x TBI interaction at 20 weeks after injury, with no main effects or interactions at 28 weeks. There was a significant increase in CD68 volume in the unstressed TBI compared to the unstressed sham mice at 5 weeks. These results are shown in Figure 4. IBA1+ cells in TBI animals appeared ameboid and had reduced processes (Figure 4A), consistent with our previous report (Hetzer et al., 2021a). At 20 weeks, there was an increase in both CD68 and IBA1+ cells, as well as a large amount of CD68 that did not overlap with Iba-1 (white arrows Figure 4A), indicating more free CD68 in injured OTs regardless of chronic stress condition. ANOVA results are reported in Table 2 with statistically significant post hoc comparisons indicated on graphs.

**Table 2.**
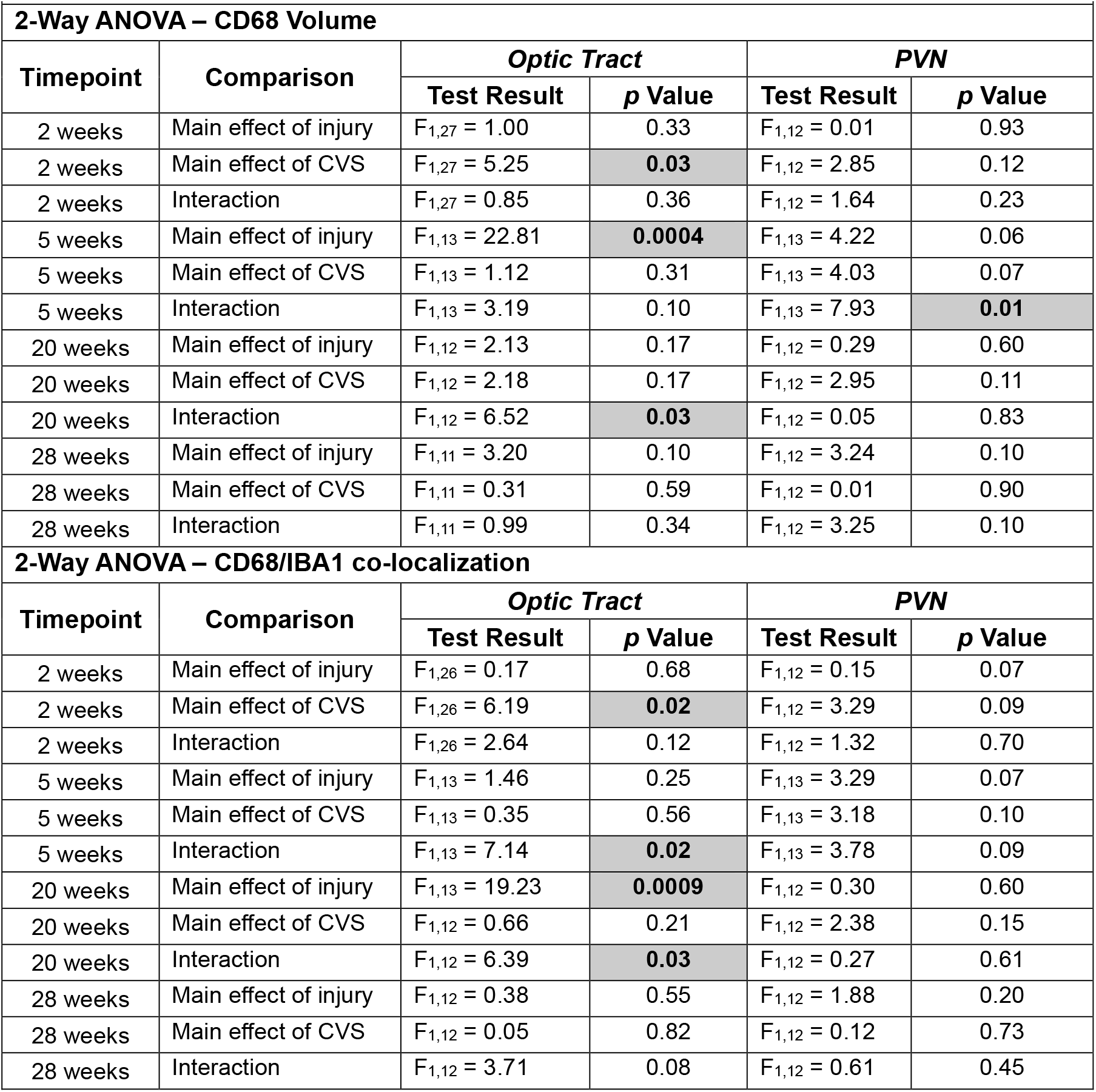
Statistical Results CD68 & Iba-1.

| **Table 2. Statistical Results CD68 & Iba-1** |  |  |  |  |  |
| --- | --- | --- | --- | --- | --- |
| **2-Way ANOVA – CD68 Volume** |  |  |  |  |  |
| **Timepoint** | **Comparison** | ***Optic Tract*** |  | ***PVN*** |  |
|  |  | **Test Result** | ***p* Value** | **Test Result** | ***p* Value** |
| 2 weeks | Main effect of injury | $F_{1,27} = 1.00$ | 0.33 | $F_{1,12} = 0.01$ | 0.93 |
| 2 weeks | Main effect of CVS | $F_{1,27} = 5.25$ | **0.03** | $F_{1,12} = 2.85$ | 0.12 |
| 2 weeks | Interaction | $F_{1,27} = 0.85$ | 0.36 | $F_{1,12} = 1.64$ | 0.23 |
| 5 weeks | Main effect of injury | $F_{1,13} = 22.81$ | **0.0004** | $F_{1,13} = 4.22$ | 0.06 |
| 5 weeks | Main effect of CVS | $F_{1,13} = 1.12$ | 0.31 | $F_{1,13} = 4.03$ | 0.07 |
| 5 weeks | Interaction | $F_{1,13} = 3.19$ | 0.10 | $F_{1,13} = 7.93$ | **0.01** |
| 20 weeks | Main effect of injury | $F_{1,12} = 2.13$ | 0.17 | $F_{1,12} = 0.29$ | 0.60 |
| 20 weeks | Main effect of CVS | $F_{1,12} = 2.18$ | 0.17 | $F_{1,12} = 2.95$ | 0.11 |
| 20 weeks | Interaction | $F_{1,12} = 6.52$ | **0.03** | $F_{1,12} = 0.05$ | 0.83 |
| 28 weeks | Main effect of injury | $F_{1,11} = 3.20$ | 0.10 | $F_{1,12} = 3.24$ | 0.10 |
| 28 weeks | Main effect of CVS | $F_{1,11} = 0.31$ | 0.59 | $F_{1,12} = 0.01$ | 0.90 |
| 28 weeks | Interaction | $F_{1,11} = 0.99$ | 0.34 | $F_{1,12} = 3.25$ | 0.10 |
| **2-Way ANOVA – CD68/IBA1 co-localization** |  |  |  |  |  |
| **Timepoint** | **Comparison** | ***Optic Tract*** |  | ***PVN*** |  |
|  |  | **Test Result** | ***p* Value** | **Test Result** | ***p* Value** |
| 2 weeks | Main effect of injury | $F_{1,26} = 0.17$ | 0.68 | $F_{1,12} = 0.15$ | 0.07 |
| 2 weeks | Main effect of CVS | $F_{1,26} = 6.19$ | **0.02** | $F_{1,12} = 3.29$ | 0.09 |
| 2 weeks | Interaction | $F_{1,26} = 2.64$ | 0.12 | $F_{1,12} = 1.32$ | 0.70 |
| 5 weeks | Main effect of injury | $F_{1,13} = 1.46$ | 0.25 | $F_{1,13} = 3.29$ | 0.07 |
| 5 weeks | Main effect of CVS | $F_{1,13} = 0.35$ | 0.56 | $F_{1,13} = 3.18$ | 0.10 |
| 5 weeks | Interaction | $F_{1,13} = 7.14$ | **0.02** | $F_{1,13} = 3.78$ | 0.09 |
| 20 weeks | Main effect of injury | $F_{1,13} = 19.23$ | **0.0009** | $F_{1,12} = 0.30$ | 0.60 |
| 20 weeks | Main effect of CVS | $F_{1,12} = 0.66$ | 0.21 | $F_{1,12} = 2.38$ | 0.15 |
| 20 weeks | Interaction | $F_{1,12} = 6.39$ | **0.03** | $F_{1,12} = 0.27$ | 0.61 |
| 28 weeks | Main effect of injury | $F_{1,12} = 0.38$ | 0.55 | $F_{1,12} = 1.88$ | 0.20 |
| 28 weeks | Main effect of CVS | $F_{1,12} = 0.05$ | 0.82 | $F_{1,12} = 0.12$ | 0.73 |
| 28 weeks | Interaction | $F_{1,12} = 3.71$ | 0.08 | $F_{1,12} = 0.61$ | 0.45 |

**Figure 4.**
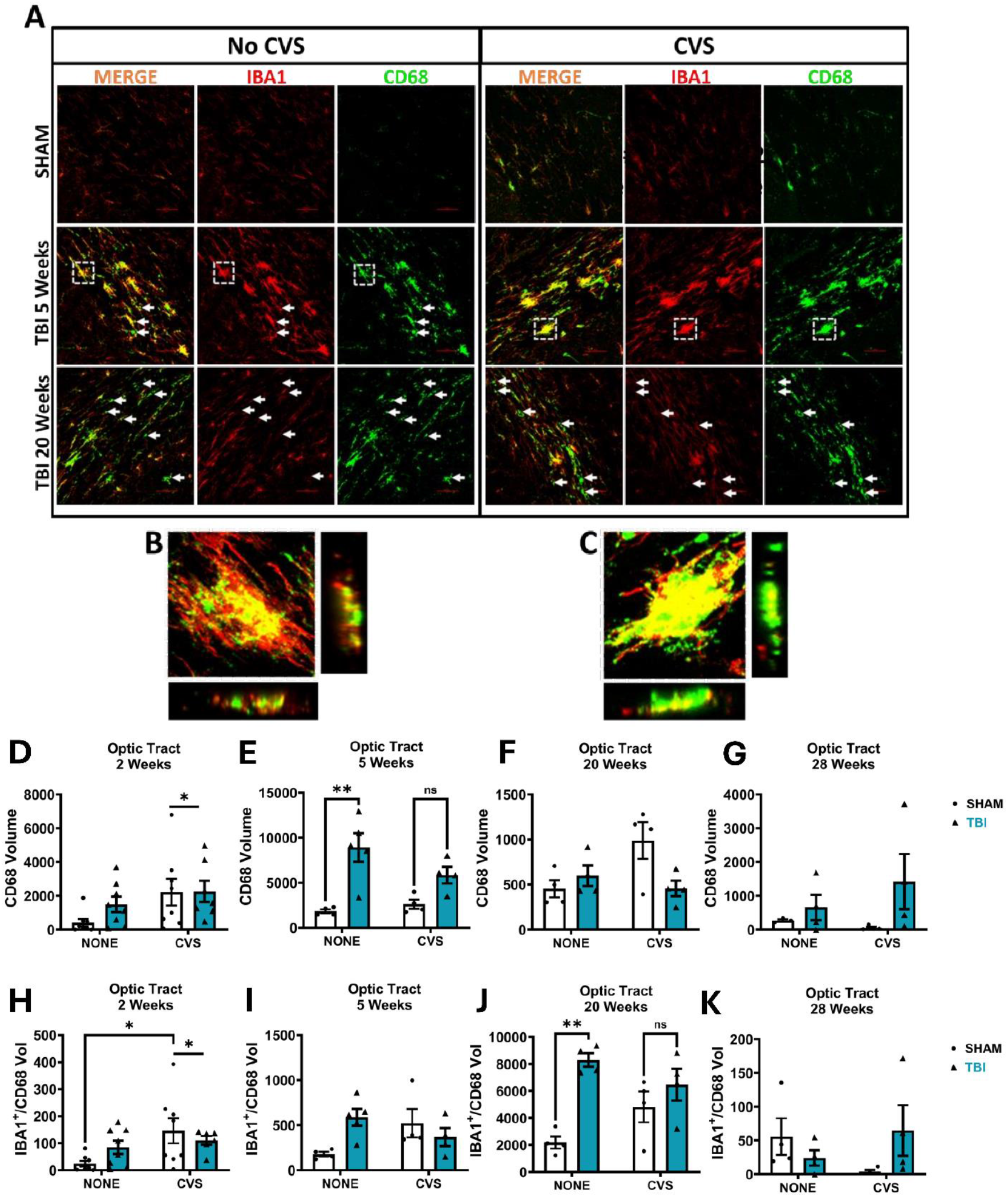
Iba-1 and CD68 immunoreactivity in optic tract. A) Representative images of Iba-1 (red), CD68 (green) and merged images in the optic tract. White arrows indicate areas positive for CD68 but not Iba-1. Higher magnification photomicrographs of individual microglia in dashed boxes are shown in (B) (TBI, no CVS) and (C) (TBI + CVS). CD68 volume in the optic tract is shown at D) 2 weeks, E) 5 weeks, F) 20 weeks, and G) 28 weeks post-injury. Volume of co-localization of Iba-1 and CD68 is shown at H) 2 weeks, I) 5 weeks, J) 20 weeks, and K) 28 weeks post-injury. There is a main effect of CVS on both CD68 volume and Iba-1/CD68 co-localization in the optic tract at 2 weeks post-injury. * p<0.05, ** p<0.01, ns: not significant.

**Figure 5:**
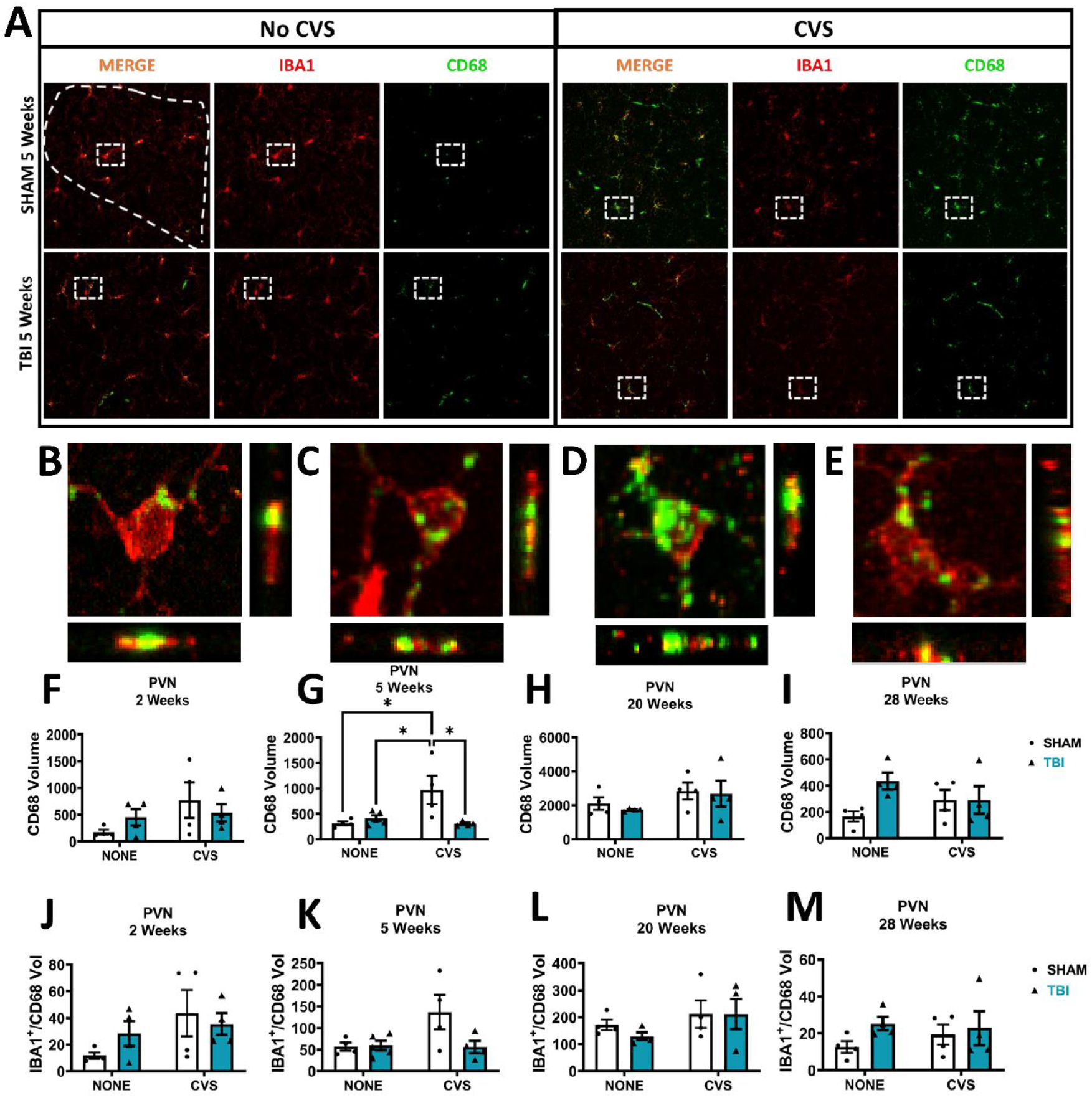
Iba-1 and CD68 immunoreactivity in PVN. A) Representative images of Iba-1 (red) and CD68 (green) immunofluorescence in the PVN, 5 weeks post-TBI. High magnification images of the area enclosed in white squares from A) with cross-sectional projections at the sides in (B) Sham, (C) TBI, (D) Sham + CVS, and (E) TBI+CVS groups at 5 weeks post-injury. (F-I) Volume of CD68 positive staining at F) 2 weeks, G) 5 weeks, H) 20 weeks, and I) 28 weeks post-injury. (J-M) Volume of pixels co-stained with Iba-1 and CD68 in the PVN at J) 2 weeks, K) 5 weeks, L) 20 weeks, and M) 28 weeks post-injury. * p<0.05.

Given the major role in regulating stress responses of the PVN, and because it receives direct optic nerve projections, we assessed Iba-1 and CD68 expression in the PVN after injury and/or CVS (Figure 5, Figure S3, and Table 2). There was a significant increase in CD68 staining in the PVN of sham + CVS mice at 5 weeks after injury compared to both TBI groups and the sham unstressed group. There was not a significant co-expression of CD68 and Iba-1 in the PVN at any time point.

We also co-stained brain tissues with antibodies against GFAP and the lysosomal marker LAMP2. In the optic tract, there was a main effect of injury on GFAP expression at 2 weeks and 28 weeks, and a CVS x TBI interaction at 20 weeks after injury (Figure 6, Table 3). For GFAP/LAMP2 co-localization, in the OT there was main effect of injury at 5 weeks, and a main effect of CVS and CVS x TBI interaction at 20 weeks. In the PVN (Figure 7), LAMP2 was not detectible in tissue from 2 weeks or 28 weeks post-injury, so it was not possible to do a co-localization analysis at these times. There were main effects of injury and CVS at 5 weeks post-injury.

**Table 3.** Statistical Results – GFAP & LAMP2.

| **Table 3. Statistical Results – GFAP & LAMP2** |  |  |  |  |  |
| --- | --- | --- | --- | --- | --- |
| **2-Way ANOVA – GFAP Volume** |  |  |  |  |  |
| **Timepoint** | **Comparison** | ***Optic Tract*** |  | ***PVN*** |  |
|  |  | **Test Result** | ***p* Value** | **Test Result** | ***p* Value** |
| 2 weeks | Main effect of injury | $F_{1,27} = 9.87$ | **0.0040** | $F_{1,12} = 4.11$ | 0.07 |
| 2 weeks | Main effect of CVS | $F_{1,27} = 3.48$ | 0.07 | $F_{1,12} = 1.36$ | 0.27 |
| 2 weeks | Interaction | $F_{1,27} = 0.01$ | 0.91 | $F_{1,12} = 1.04$ | 0.33 |
| 5 weeks | Main effect of injury | $F_{1,13} = 5.42$ | 0.21 | $F_{1,13} = 2.99$ | 0.11 |
| 5 weeks | Main effect of CVS | $F_{1,13} = 0.74$ | 0.40 | $F_{1,13} = 0.36$ | 0.56 |
| 5 weeks | Interaction | $F_{1,13} = 1.77$ | 0.20 | $F_{1,13} = 0.04$ | 0.85 |
| 20 weeks | Main effect of injury | $F_{1,11} = 2.44$ | 0.14 | $F_{1,12} = 0.02$ | 0.89 |
| 20 weeks | Main effect of CVS | $F_{1,11} = 1.60$ | 0.23 | $F_{1,12} = 0.05$ | 0.83 |
| 20 weeks | Interaction | $F_{1,11} = 9.64$ | **0.01** | $F_{1,12} = 6.02$ | **0.03** |
| 28 weeks | Main effect of injury | $F_{1,12} = 8.98$ | **0.01** | $F_{1,12} = 0.07$ | 0.80 |
| 28 weeks | Main effect of CVS | $F_{1,12} = 1.81$ | 0.20 | $F_{1,12} = 2.96$ | 0.11 |
| 28 weeks | Interaction | $F_{1,12} = 0.99$ | 0.34 | $F_{1,12} = 0.19$ | 0.67 |
| **2-Way ANOVA – GFAP/LAMP2 co-localization** |  |  |  |  |  |
| **Timepoint** | **Comparison** | ***Optic Tract*** |  | ***PVN*** |  |
|  |  | **Test Result** | ***p* Value** | **Test Result** | ***p* Value** |
| 2 weeks | Main effect of injury | $F_{1,25} = 2.94$ | 0.10 | No LAMP2 | - |
| 2 weeks | Main effect of CVS | $F_{1,25} = 0.66$ | 0.44 | No LAMP2 | - |
| 2 weeks | Interaction | $F_{1,25} = 0.62$ | 0.44 | No LAMP2 | - |
| 5 weeks | Main effect of injury | $F_{1,13} = 10.94$ | **0.006** | $F_{1,13} = 4.85$ | **0.046** |
| 5 weeks | Main effect of CVS | $F_{1,13} = 0.002$ | 0.97 | $F_{1,13} = 7.16$ | **0.02** |
| 5 weeks | Interaction | $F_{1,13} = 0.21$ | 0.65 | $F_{1,13} = 3.54$ | 0.08 |
| 20 weeks | Main effect of injury | $F_{1,11} = 0.03$ | 0.87 | $F_{1,12} = 3.25$ | 0.1 |
| 20 weeks | Main effect of CVS | $F_{1,11} = 5.15$ | **0.04** | $F_{1,12} = 1.26$ | 0.28 |
| 20 weeks | Interaction | $F_{1,11} = 6.86$ | **0.02** | $F_{1,12} = 1.51$ | 0.24 |
| 28 weeks | Main effect of injury | $F_{1,12} = 4.53$ | 0.055 | No LAMP2 | - |
| 28 weeks | Main effect of CVS | $F_{1,12} = 0.23$ | 0.64 | No LAMP2 | - |
| 28 weeks | Interaction | $F_{1,12} = 0.11$ | 0.74 | No LAMP2 | - |

**Figure 6.**
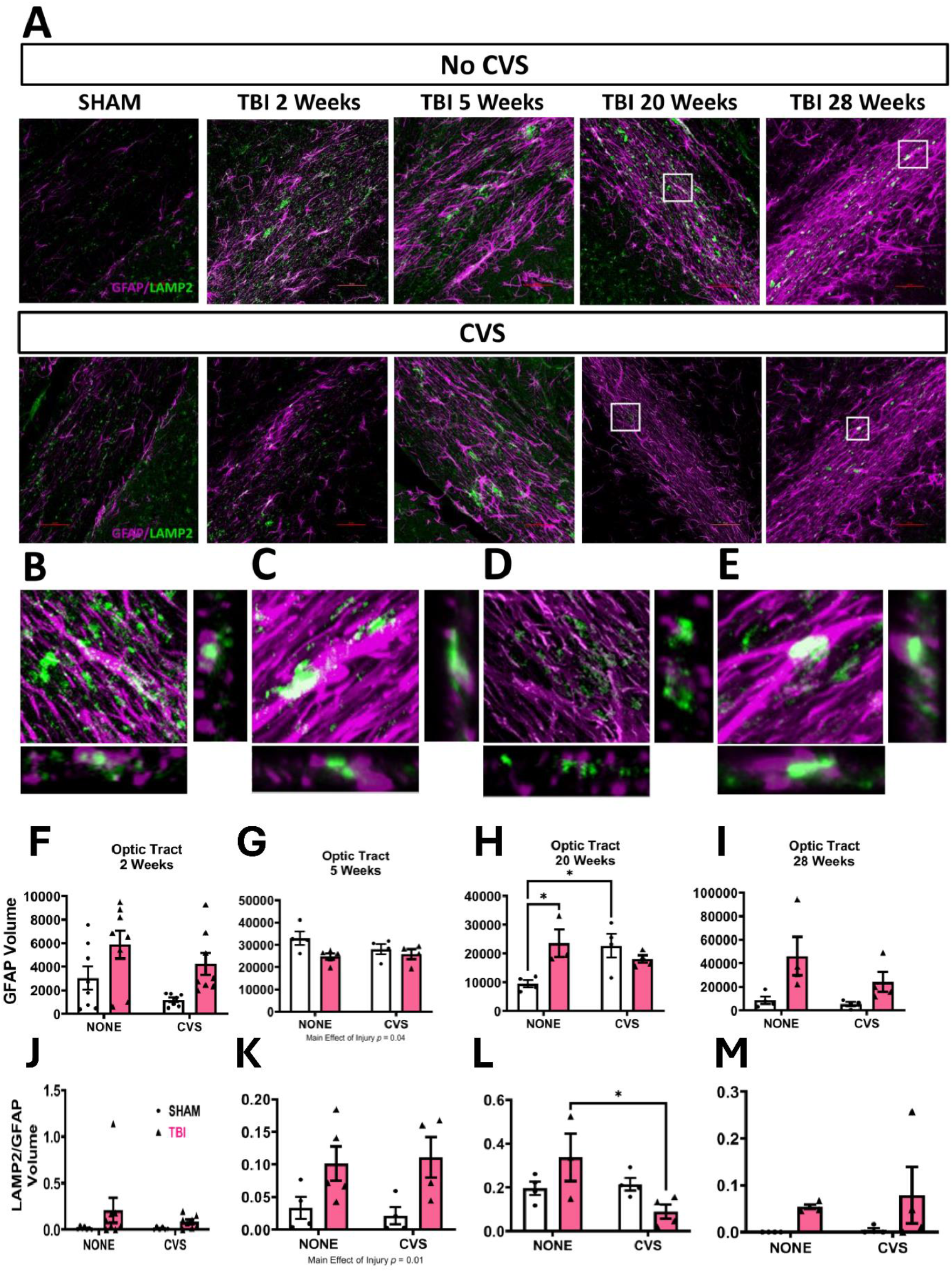
GFAP and LAMP2 Immunoreactivity in the optic tract. (A) Representative images of GFAP (purple) and LAMP2 (green) co-staining of the optic tract. Also shown are higher-magnification photomicrographs from the regions outlined in white in part A) from B) 20-week TBI, C) 28-week TBI, D) 20-week TBI/CVS and E) 28-week TBI/CVS. Volume of GFAP expression in the optic tract at F) 2 weeks, G) 5 weeks, H) 20 weeks, or I) 28 weeks post-injury, and of GFAP and LAMP2 co-expression at J) 2 weeks, K) 5 weeks, L) 20 weeks, and M) 28 weeks are shown. * p<0.05.

**Figure 7.**
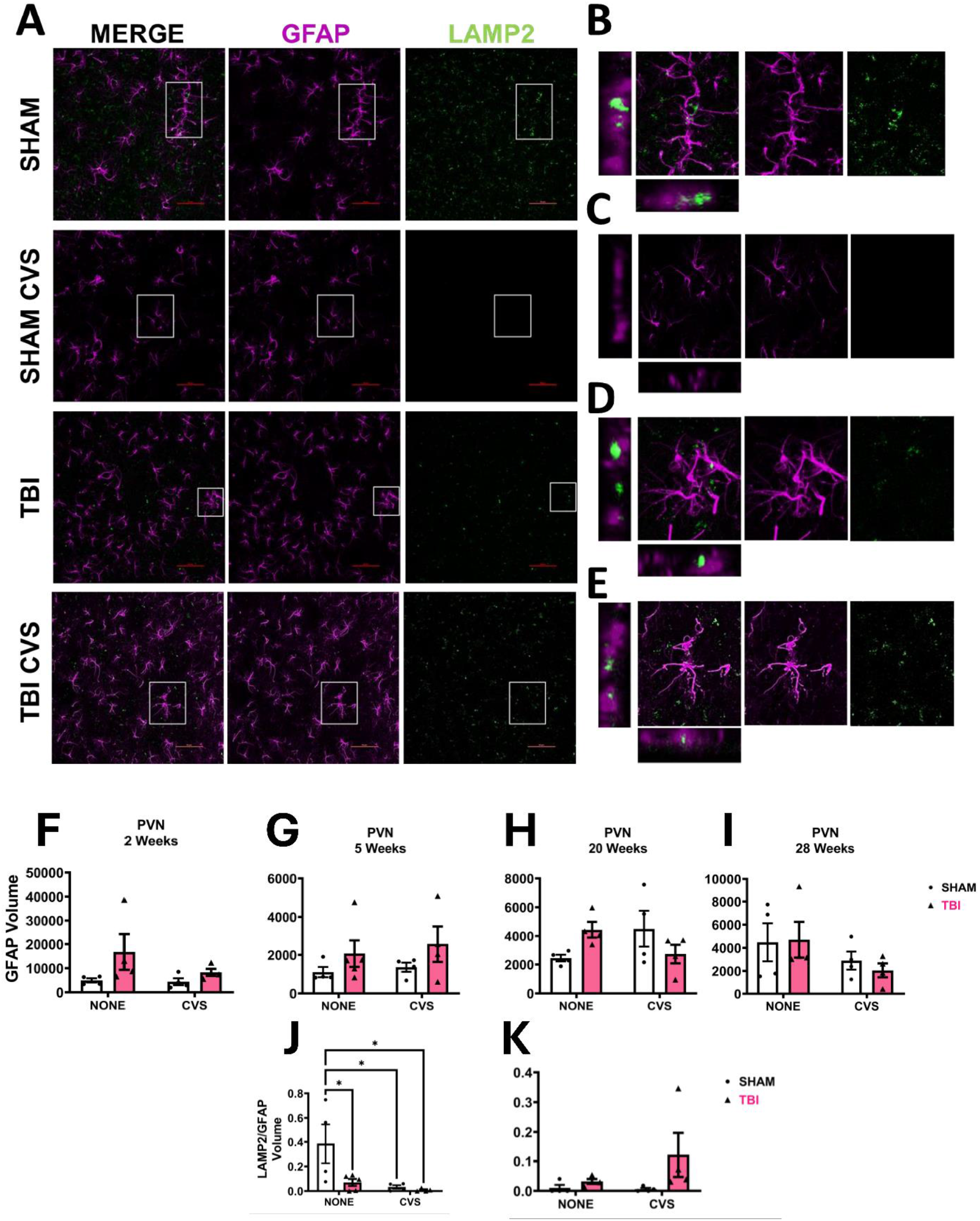
GFAP and LAMP2 Immunoreactivity in the PVN. (A) Representative images of LAMP2 (green) and GFAP (purple) co-staining of the PVN at 20 weeks post-injury. (B-E) High-resolution photomicrographs of the region in white boxes in part A, with lateral projections shown beside the merged images. (F-I) Shows GFAP volume over four timepoints within the PVN. No significance was found within this region. (J-K) Volume of GFAP and LAMP2 co-expression in the PVN at J) 5 weeks and K) 20 weeks after TBI. There was very little LAMP2 expression detectible in the PVN overall, and none in the 2 week and 28 week cohorts, which are thus not shown. * p<0.05.

## Discussion

TBI is a significant health issue in adolescents, often leading to long-term disability. Importantly, early treatment of patients who are hospitalized after TBI often involves multiple stressful events necessary in the ICU setting (Krampe et al., 2021). Although this treatment may improve survival and reduce secondary injury (Kochanek et al., 2019), it is possible that the chronic stress of the treatment could have negative consequences for recovery. Chronic stress can impair cognitive function (Kim and Kim, 2023) and is linked to increased risk of neurodegenerative conditions (Peña-Bautista et al., 2020). Given these potentially negative consequences of chronic stress, we hypothesized that chronic stress administered early after TBI leads to worse chronic TBI outcomes in an adolescent murine head trauma model. We found instead that the early effects of CVS on tissue pathology associated with head trauma were mild, and, surprisingly, that there is potentially a delayed neuroprotective effect of CVS on measures of axonal degeneration in the optic tract and in its major projection targets.

Specifically, post-injury CVS led to absence of axonal degeneration as measured by FJ-B staining 20-weeks post-injury. This time point was also associated with changes in immunohistological markers of astrocyte and microglial activation and phagocytosis, which suggest a possible mechanism for the protective effect. As expected, there was an early effect of TBI on weight gain. There were also persistent effects of CVS and TBI on overall weight gain, with CVS and TBI groups having slightly less long-term weight gain than control animals. This finding supports chronic effects of both CVS and TBI on these mice. There was not an effect of TBI or CVS on forced swim test or novel object recognition test results, suggesting recovery of these measures by the end of the CVS period.

We previously reported that TBI leads to optic nerve injury and axonal degeneration in the optic tracts and multiple optic tract projection targets, most prominently including the lateral geniculate nucleus of the thalamus and superior colliculus (Evanson et al., 2018). Optic tract degeneration persisted through 150 days post-injury in those studies (Hetzer et al., 2021a). The current results confirm and extend those previous findings. We found in this study that CVS led to elimination of FJ-B staining in the optic tracts 20 weeks after TBI, followed by return of FJ-B staining 28 weeks after injury. Because FJ-B stains degenerating neurons and axons (Schmued and Hopkins, 2000), our results suggest the presence of degenerating axons in all of the TBI conditions examined except at 20 weeks after injury in the CVS/TBI group.

After traumatic axon breakage, distal axon segments degenerate, often over long periods of time, in a process known as Wallerian Degeneration (Coleman and Höke, 2020). In this process, distal axon segments degrade and proximal segments retract to the cell somata. Microglia, astrocytes, and invading immune cells contribute to clearing cellular and distal axonal debris during this process (Neumann et al., 2009; Yu et al., 2021). Particularly for axonal debris, this clearing process can occur over a long period of time, with degenerating axons sometimes persisting for months or longer (Johnson et al., 2013). From the FJ-B staining alone in the current studies, it is not clear whether these degenerating axons are present because they have not yet been cleared or because there is ongoing degeneration occurring. However, the fact that FJ-B stained material in the optic tract is absent at 20-weeks post-injury in TBI mice subjected to CVS, but reappears at 28-weeks post-TBI suggests that ongoing degeneration is likely occurring in these mice. This interpretation is supported by findings from other groups that optic nerve stretch injury leads to ongoing optic nerve axon loss for at least 12 weeks after injury (Maxwell et al., 2015).

Consistent with reported roles for microglia and astrocytes in clearing cellular debris after injury (as noted above), we found immunohistologic evidence of active phagocytosis occurring via both microglia and astrocytes in optic tracts, based on co-localization of Iba-1 and CD68 as well as GFAP and LAMP2. CD68 is a membrane-associated lysosomal protein that is enriched in phagocytically active microglia (Yang et al., 2022), while Iba-1 is used as an immunocytochemical marker for microglia and macrophages (Ahmed et al., 2007). Co-localization of these markers, therefore, is a marker for active phagocytosis by microglia (Hendrickx et al., 2017). Similarly, LAMP2 is a lysosome-associated protein, and its expression in astrocytes is increased in reactive astrocytes (Fu et al., 2023). It is also associated with astrocyte phagocytic activity (Morizawa et al., 2017). Our results suggest that TBI may lead to increased phagocytic activity in the optic tracts, which was most prominently detected after 5 weeks post-injury.

In this study, microglial activation was significantly elevated by 5 weeks after TBI, as evidenced by increased CD68 co-expression (Figure 4E). Microglial phagocytosis, marked by elevated Iba-1/CD68 co-expression, was elevated in TBI mice 20 weeks after injury, but not in TBI/CVS mice, compared to their controls. This change likely starts earlier than 20 weeks, as evidenced by a significant interaction in the 5 week Iba-1/CD68 data. At 2 weeks after injury, CVS alone increases both CD68 and Iba-1/CD68 co-localization (Figure 4D, H). These results suggest that CVS causes biphasic changes in neuro-inflammation, with an increase early after CVS is completed, but with a delayed long-term decrease in neuro-inflammation in TBI mice.

Chronic stress acutely increases blood-brain barrier permeability and oxidative stress (Tagliari et al., 2010b; Lehmann et al., 2018), while reducing brain energy metabolism (Tagliari et al., 2010a). Both acute and chronic stress can cause release of pro-inflammatory cytokines and induce microglial activation (Walker et al., 2013). Chronic stress also leads to increased phagocytic activity of microglia (Kokkosis et al., 2024). These findings are consistent with our findings of increased CD68/Iba-1 co-expression in CVS mice by 2 weeks after injury. However, our results also indicate that CVS effects are not solely harmful, because of the delayed protective effect of CVS on axonal degeneration seen. This is consistent with the roles of neuro-inflammation on long-term brain injury outcomes, which includes both damaging and protective or reparative roles (Theus, 2024). Similarly, microglia can have both damaging, pro-inflammatory function, and also homeostatic or reparative activity (Butovsky and Weiner, 2018). Chronic stress can also have an anti-inflammatory effect on microglial responses (Smith et al., 2016), which could be a potential mechanism for the current findings.

It is important to note that mice in this study were 6 weeks old at the onset of CVS, and CVS continued until 8 weeks of age. Six weeks of age is generally considered to represent adolescence in a mouse, while 8 weeks is considered adult (Semple et al., 2016). This may be an important factor in the results of these studies, because adolescence is a time of increased plasticity in response to stress (Sisk and Gee, 2022), and CVS exposure can have different long-term effects depending on whether stress is administered during adolescence or adulthood (Jankord et al., 2011; Cotella et al., 2019). In fact, chronic stress in adolescence tends to have effects on behavior and brain structure persisting into adulthood, while such changes are likely to be transient when chronic stress is experienced in adulthood (Hollis et al., 2013). Interestingly, chronic unpredictable stress experienced during adolescence is protective against cognitive effects of TBI sustained in adulthood in a controlled cortical impact model (de la Tremblaye et al., 2021). Our findings may hint at a mechanism for this effect (changes in neuroinflammation in axon tracts), although the order and timing of the chronic stress exposure and TBI are different. Future work should address the importance of these differences in approach.

The paraventricular nucleus of the hypothalamus (PVN) is critical in integrating brain stress responses and regulating hypothalamus-pituitary-adrenal (HPA) axis responses to stress. It also plays a significant role in chronic stress adaptation (Herman and Tasker, 2016). The PVN receives direct projections from the optic nerve (Schaechter and Sadun, 1985; Youngstrom et al., 1991) as well as from the lateral geniculate nucleus (Mikkelsen, 1990). Although the function of these projections has not been explored, it is potentially related to the fact that PVN activity is regulated in a circadian fashion (Jones et al., 2021). Because the PVN receives input from the retina and plays a central role in chronic stress adaptation, we assessed the PVN in the current studies. We found that CD68 volume was increased in the PVN 5 weeks after injury (3 weeks after CVS), in the CVS-only control group. This increase in CD68 volume is consistent with previous reports that CVS causes neuro-inflammation in the PVN, with an associated increase in CD68-expressing macrophages (Lehmann et al., 2016). Activated microglia in the PVN likely lead to increased neuronal activity in the PVN (Han et al., 2021). Thus, CVS could potentially be exerting some effects in this system via pro-inflammatory effects of CVS on the PVN. On the other hand, TBI leads to HPA axis hyper-activation acutely after TBI (Shohami et al., 1995; McCullers et al., 2002), and can cause hypo-activity subacutely after TBI (Taylor et al., 2006). Thus, these 5-week findings may be consistent with changes in HPA axis activity at this time point (increased from CVS, and balanced/decreased by TBI combined with CVS). Further study will be needed to better understand this relationship between TBI, CVS, and HPA axis activity.

The current studies focused on brain histologic markers of injury because at the time these studies were initiated, it was not yet clear that the model used primarily affected the optic tracts. Thus, our results do not include significant functional measures related to optic nerve integrity. Future work will also need to more fully assess phagocytic activity of astrocytes and microglia, as well as myelin integrity of the optic tracts at chronic post-injury time points. In addition, we do not have humoral measures of HPA axis activity during this study due to the focus on brain measures of injury and neuronal/axonal degeneration. Thus, future studies will need to include functional measures of optic nerve activity/integrity, better characterization of phagocytic activity by microglia and astrocytes, and assessment of humoral measures of HPA axis activity.

## Conclusion

In conclusion, chronic variable stress after TBI leads to lasting, but not permanent, effects on axon injury and degeneration in the optic tracts, with a protective effect on chronic optic tract axonal degeneration. There is a peak effect around 20 weeks after TBI, or 18 weeks after the end of CVS, with suspected lessening of the protective effect after this time. CVS independently increases markers associated with microglial phagocytosis early after injury. TBI alone leads to subacute and chronic increase in markers of microglial activation 20 weeks after injury, but this effect is reduced by CVS. Thus, it is possible that CVS exerts its protective effect on optic tract axonal degeneration by altering microglial activity.

## Statements and Declarations

The authors declare no conflicts of interest.

Animal work was approved by the University of Cincinnati, IACUC protocol # 17-04-03-01.

## Acknowledgments

This work was funded by grants from the National Institutes of Health (HD001097 to NKE and NS007453 to SMH) and from the Department of Defense Vision Research Program (W81XWH-22-1-1075 to NKE), and by a Procter Scholar Award from Cincinnati Children’s Hospital to NKE.

