## Supplementary information for "Delayed protective effect of chronic variable stress on optic tract axonal degeneration after experimental TBI"

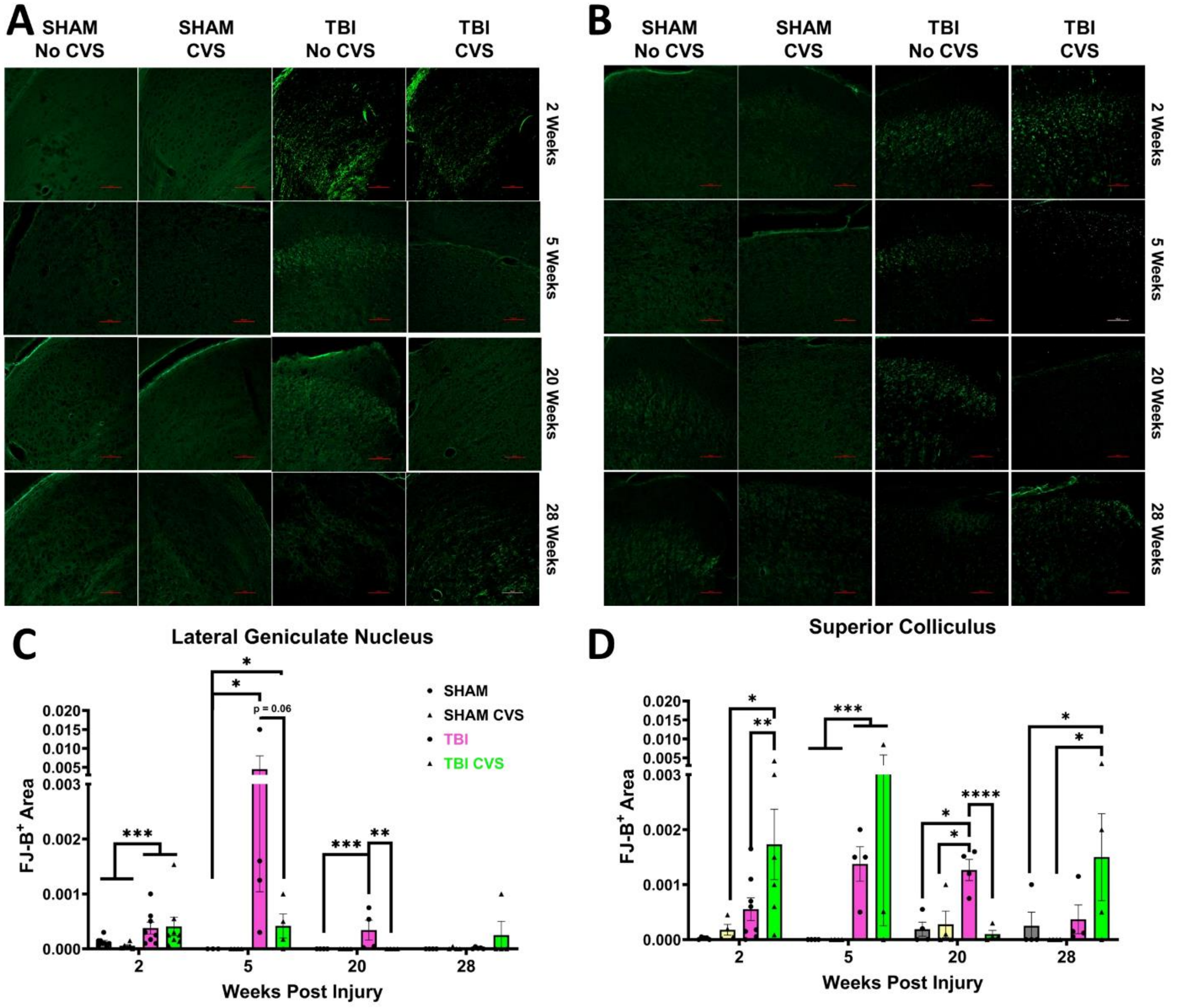


**Supplementary Figure 1. FJ-B staining in LGN and SC.** Representative images of FJ-B staining in A) LGN and B) SC at 2, 5, 10, and 28 weeks post-injury. (C) Quantification of FJ-B staining in C) LGN and D) SC. * p<0.05, ** p<0.01, ***p<0.001, ****P<0.0001.

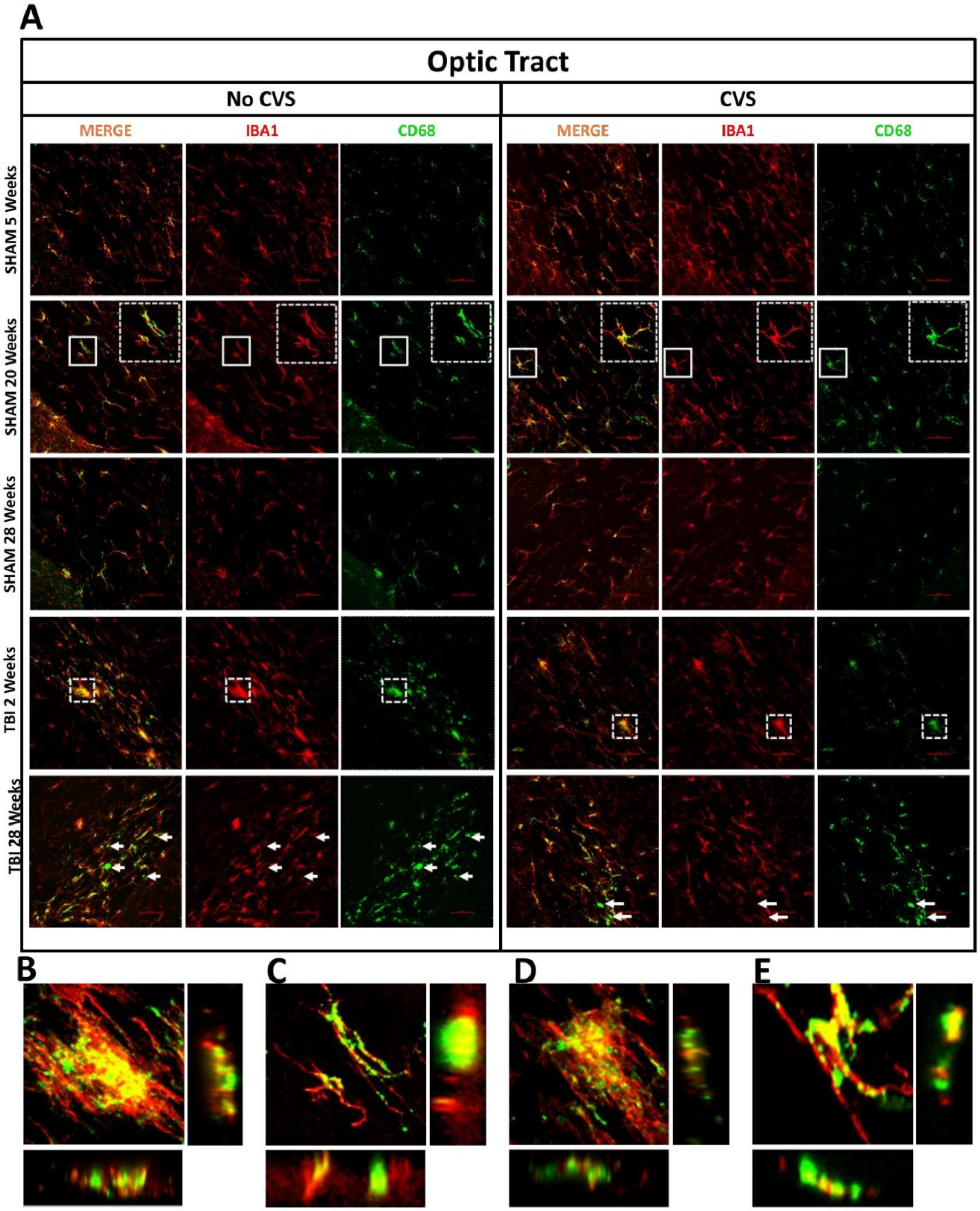


**Supplementary Figure 2. Additional representative images for Iba-1 and CD68 co-staining of optic tracts.** A) Images from the remainder of time points not shown in Figure 4. (B-E) Higher resolution photomicrographs of the areas shown in boxes from part A, from B) 2 week TBI group, C) 20 week sham, D) 2 week TBI/CVS, and E) 20 week sham/CVS groups. Lateral projections of the z-stacks are shown beside and below each image. White arrows indicate areas positive for CD68 but not Iba-1.

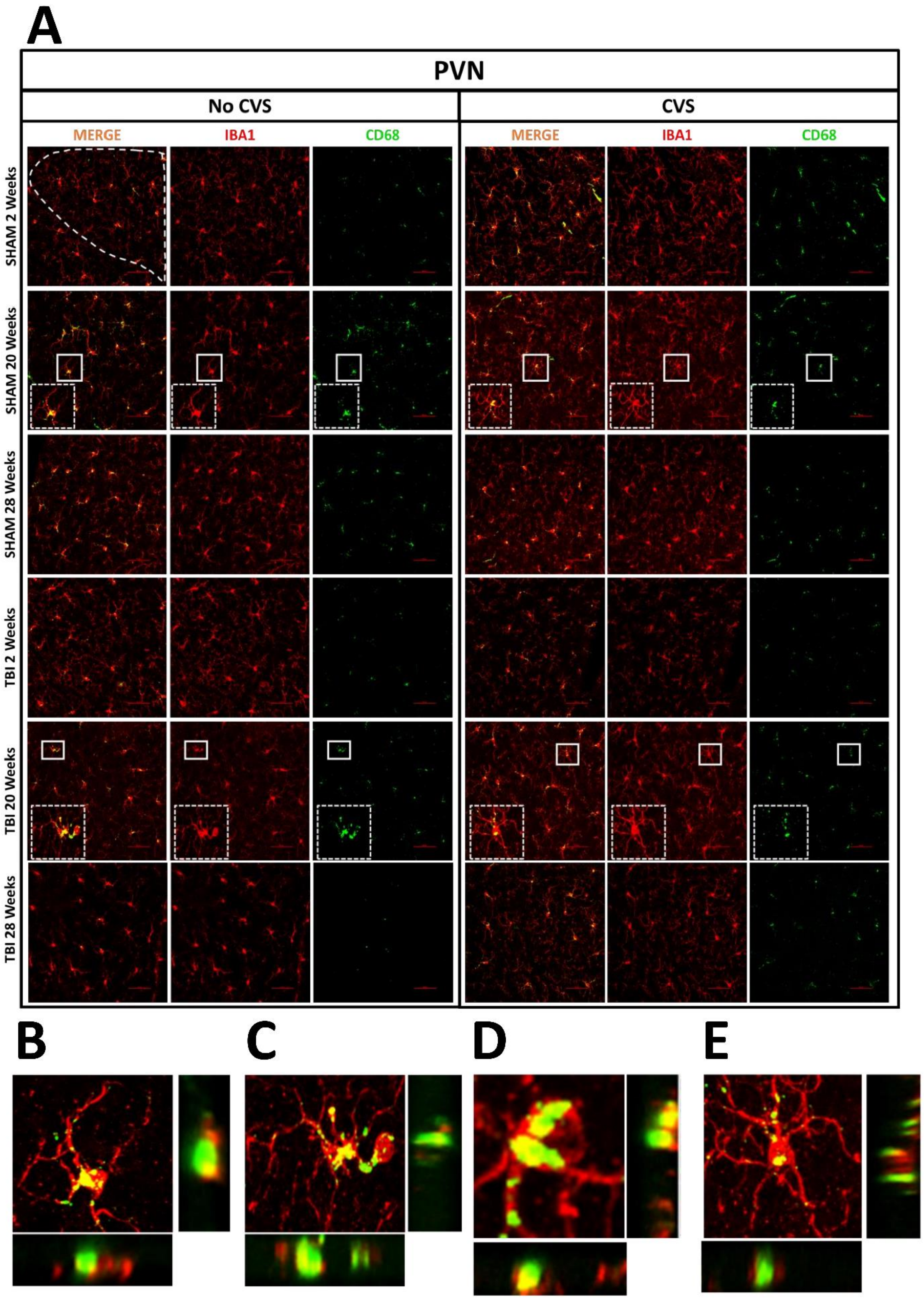

A
PVN
No CVS
CVS
MERGE
IBA1
CD68
MERGE
IBA1
CD68
SHAM 2 Weeks
SHAM 20 Weeks
SHAM 28 Weeks
TBI 2 Weeks
TBI 20 Weeks
TBI 28 Weeks
B
C
D
E

**Supplementary Figure 3. Additional representative images for IBA1 and CD68 co-staining in PVN.** (A) Representative images of the PVN showing Iba-1 (red) and CD68 (green) staining as labeled, for groups not shown in Figure 5. (B-E) Higher magnification photomicrographs of areas outlined in white boxes in part A, with lateral projection views beside and below each image. Images are from B) Sham 20 weeks, C) TBI 20 weeks, D) Sham + CVS 20 weeks, and E) TBI+CVS 20 weeks.

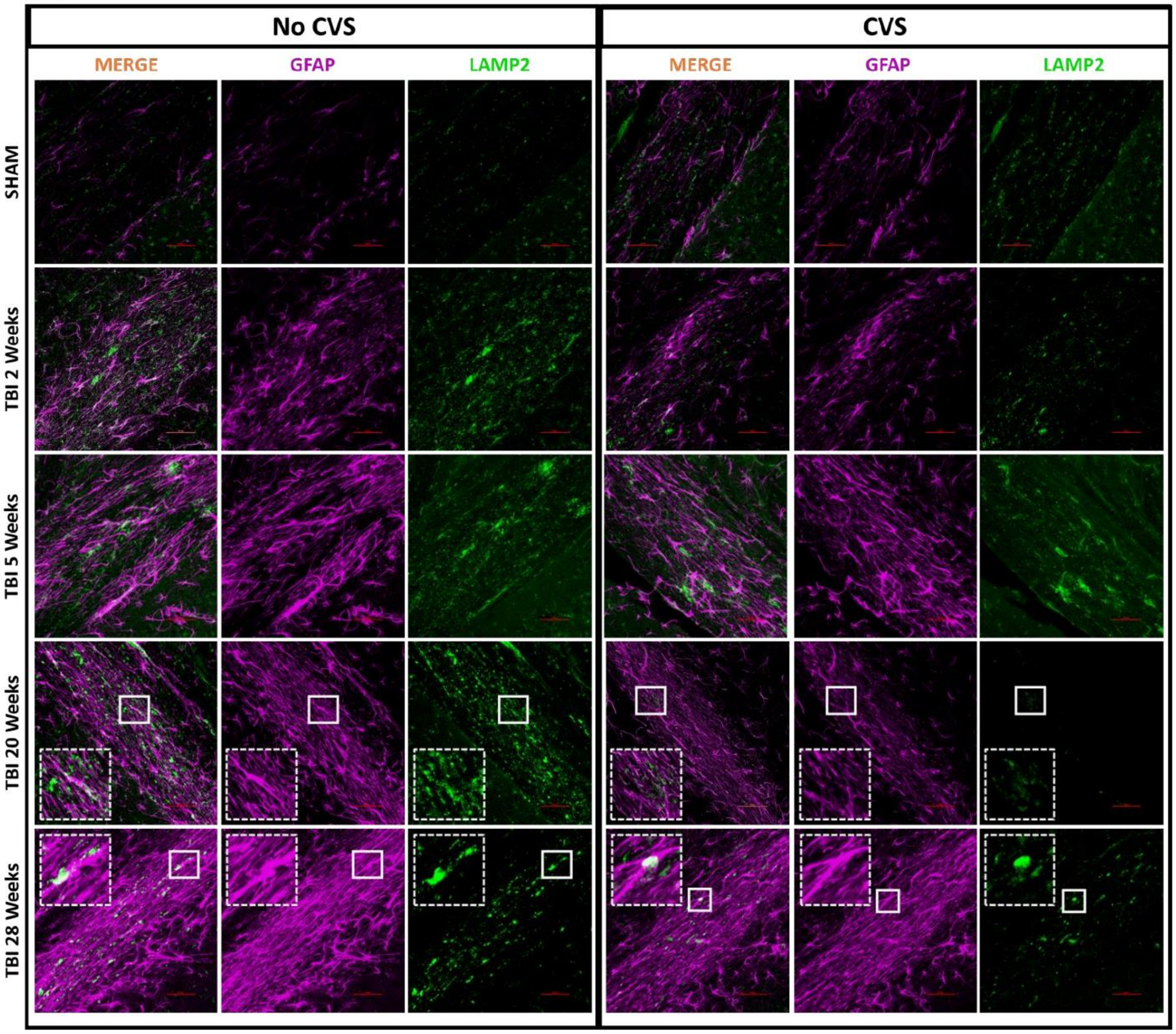


**Supplementary Figure 4.** Additional representative images of LAMP2-(green) and GFAP-(purple) immunofluorescence in the optic tract. Sham group images are from the 20 week time point. Inset images in dashed boxes are higher-magnification pictures of the areas in solid white boxes.

**Supplementary Table S1. Weight Test Statistics**

### Mixed effects analysis

| Comparison | Test Result | *p* Value |
|---|---|---|
| Injury | $F_{1, 140} = 34.82$ | **<0.0001** |
| CVS | $F_{1, 140} = 11.02$ | **0.001** |
| Time x Injury | $F_{11, 807} = 4.221$ | **<0.0001** |
| Time x CVS | $F_{11, 807} = 2.902$ | **0.0009** |
| Injury x CVS | $F_{1, 140} = 0.6095$ | 0.43 |
| Time x Injury x CVS | $F_{11, 807} = 1.529$ | 0.11 |

### Significant Fisher's LSD Post-hoc Tests

| Comparison Day | Comparison Groups | *p* Value | Comparison Day | Comparison Groups | *p* Value |
|---|---|---|---|---|---|
| Day 1 | SHAM vs SHAM CVS | 0.02 | Day 11 | SHAM vs SHAM CVS | 0.03 |
| Day 1 | SHAM vs TBI | <0.0001 | Day 11 | SHAM vs TBI | 0.04 |
| Day 1 | SHAM vs TBI CVS | 0.0009 | Day 11 | SHAM vs TBI CVS | <0.0001 |
| Day 1 | SHAM CVS vs TBI | <0.0001 | Day 11 | SHAM CVS vs TBI CVS | 0.001 |
| Day 1 | SHAM CVS vs TBI CVS | 0.0001 | Day 11 | TBI vs TBI CVS | 0.0003 |
| Day 3 | SHAM vs TBI | 0.0006 | Day 14 | SHAM vs SHAM CVS | 0.008 |
| Day 3 | SHAM vs TBI CVS | <0.0001 | Day 14 | SHAM vs TBI | 0.006 |
| Day 3 | SHAM CVS vs TBI CVS | 0.002 | Day 14 | SHAM vs TBI CVS | <0.0001 |
| Day 5 | SHAM vs TBI CVS | 0.01 | Day 14 | SHAM CVS vs TBI CVS | 0.0007 |
| Day 7 | SHAM vs TBI | 0.04 | Day 14 | TBI vs TBI CVS | 0.001 |
| Day 9 | SHAM vs TBI | 0.03 | Day 21 | SHAM vs SHAM CVS | 0.03 |
| Day 9 | SHAM vs TBI CVS | <0.0001 | Day 21 | SHAM vs TBI | 0.01 |
| Day 9 | SHAM CVS vs TBI CVS | 0.004 | Day 21 | SHAM vs TBI CVS | 0.003 |
| Day 9 | TBI vs TBI CVS | 0.03 | Day 28 | SHAM vs TBI CVS | 0.04 |

| **Supplementary Table S2. FJ-B 2-Way ANOVA – LGN & SC** | | | | | |
|---|---|---|---|---|---|
| **Timepoint** | **Comparison** | ***LGN*** | | ***SC*** | |
| | | **Test Result** | ***p* Value** | **Test Result** | ***p* Value** |
| 2 Weeks | Main effect of injury | $F_{1,26}$ = 7.58 | **0.01** | $F_{1,19}$ = 7.52 | **0.01** |
| 2 Weeks | Main effect of stress | $F_{1,26}$ = 0.03 | 0.85 | $F_{1,19}$ = 3.07 | 0.10 |
| 2 Weeks | Interaction | $F_{1,26}$ = 0.21 | 0.65 | $F_{1,19}$ = 1.81 | 0.19 |
| 5 Weeks | Main effect of injury | $F_{1,12}$ = 0.39 | 0.54 | $F_{1,11}$ = 2.58 | 0.14 |
| 5 Weeks | Main effect of stress | $F_{1,12}$ = 0.13 | 0.72 | $F_{1,11}$ = 0.92 | 0.36 |
| 5 Weeks | Interaction | $F_{1,12}$ = 0.12 | 0.73 | $F_{1,11}$ = 1.42 | 0.26 |
| 20 Weeks | Main effect of injury | $F_{1,12}$ = 1 .18 | 0.30 | $F_{1,12}$ = 6.94 | **0.02** |
| 20 Weeks | Main effect of stress | $F_{1,12}$ = 0.85 | 0.38 | $F_{1,12}$ = 9.94 | **0.01** |
| 20 Weeks | Interaction | $F_{1,12}$ = 0.001 | 0.97 | $F_{1,12}$ = 13.43 | **0.003** |
| 28 Weeks | Main effect of injury | $F_{1,13}$ = 1.40 | 0.26 | $F_{1,12}$ = 1.01 | 0.33 |
| 28 Weeks | Main effect of stress | $F_{1,13}$ = 4.22 | 0.06 | $F_{1,12}$ = 1.42 | 0.26 |
| 28 Weeks | Interaction | $F_{1,13}$ = 3.06 | 0.26 | $F_{1,12}$ = 1.82 | 0.20 |
